# Explicit derivation of the standard fitting technique for two-state protein transitions using differential scanning calorimetry data

**DOI:** 10.1101/2024.11.01.621328

**Authors:** Tadeas Priklopil

## Abstract

This paper has three main objectives. First, we aim to derive from first principles a linear heat capacity model for proteins exhibiting two-state conformational transition dynamics, which includes, as a special case, the constant heat capacity model frequently referenced in the literature. Second, we present a protocol for inferring the Gibbs free energy of protein folding by fitting the linear heat capacity model to differential scanning calorimetry data. Third, we demonstrate this protocol using Mathematica software, applying it to real datasets and comparing the linear and constant heat capacity models. While most of our derivations are well established in the literature, we provide a step-by-step bridge from mathematical modeling to application, addressing a practical gap in existing resources that benefits researchers.

## 1 Introduction

Differential scanning calorimetry (DSC) is a reliable and widely used method for assessing the thermodynamic stability profiles of proteins (see, for example, Becktel and Schellman, 1987; Hemminger and Sarge, 1991; Freire, 1995; McCrary et al., 1996; Fersht, 1999; Höhne et al., 2003; Privalov, 2012; Pucci and Rooman, 2014; Seelig and Schönfeld, 2016; Barrick, 2018; Gibson et al., 2022; Yeritsyan and Badasyan, 2024a,b). Originally introduced by Watson and O’Neill (1962) and further developed by Privalov et al. (1964), DSC enables precise measurement of heat capacity changes during thermal transitions in proteins. By capturing the unfolding process, DSC allows researchers to estimate key thermodynamic parameters that define a protein’s stability profile. These profiles reveal protein stability across temperatures, providing insights applicable in fields such as protein engineering for industrial applications (e.g., Frokjaer and Otzen, 2005; Ebrahimi and Samanta, 2023) and in population genetics to model adaptive evolution of biophysical properties in natural populations (Berger et al., 2021, Priklopil et al., 2024).

Despite its widespread use, complete procedures for deriving mathematical models and fitting DSC data to them are often incomplete or scattered across various sources (for example, Biltonen et al., 1978; Freire, 1995; McCrary et al., 1996; Spink, 2008; Durowoju et al., 2017; Barrick, 2018; Seelig and Schönfeld, 2016; Gibson et al., 2022; Yeritsyan and Badasyan, 2024a,b). In this paper, we address this gap by providing a detailed derivation of a linear heat capacity (LHC) model for proteins exhibiting two-state transition dynamics (Section 2), which effectively captures the folding dynamics of many small, single-domain proteins (Privalov, 2012). This LHC model extends the constant heat capacity (CHC) model, which assumes the heat capacities of folded and unfolded proteins are linear with temperature and parallel (i.e., having the same slope) (Barrick, 2018), by allowing for non-parallel heat capacities. This adjustment better reflects observations for some two-state proteins with minor structural variations (Privalov and Dragan, 2007; LiCata and Liu, 2011). We then introduce a protocol for fitting this model to DSC data to infer the Gibbs free energy of protein folding (Section 3) and demonstrate the procedure using real data (see online supplement, Priklopil, 2024). While not entirely novel, our aim is to provide a comprehensive resource to help researchers understand and effectively apply this methodology.

## 2 Linear heat capacity model for two-state proteins

Our derivation of the LHC model is based on applying reaction thermodynamics to equilibrium conformational transitions of proteins (Barrick, 2018).

### Equilibrium conformational transitions of two-state proteins

We will consider a class of proteins that can to a good approximation be in just one of two thermodynamical states, a folded F (or native) state and an unfolded U (or denaturated) state, between which they can transition. We will thus be considering a reversible two-state reaction

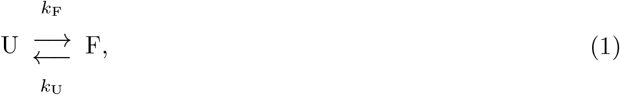

where *k*_*U*_, *k*_*F*_ are temperature-dependent rates at which an unfolded protein transitions to the folded state and a folded protein transitions to the unfolded state, respectively. Supposing that at any given time the total number of proteins 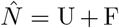 is large and constant (and where we abuse the notation by denoting with U, F the concentration as well as the state of the protein), the folding-unfolding dynamics can be expressed as 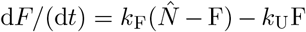. Supposing that the folding-unfolding thermodynamics is at its equilibrium (the rate of heating is a much slower process than folding-unfolding dynamics), which can be solved from d*F/*(d*t*) = 0, and which can be usefully expressed as

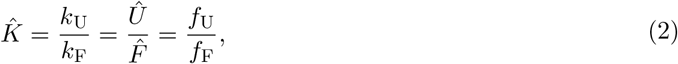

where 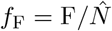 and 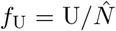, with *f*_F_ + *f*_U_ = 1, are the frequencies of folded and unfolded proteins at the equilibrium, respectively. We emphasise that since the rates *k*_F_, *k*_U_ are temperature dependent, so is the equilibrium 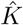. At equilibrium, the Gibbs free energy difference between the folded and unfolded states is given by

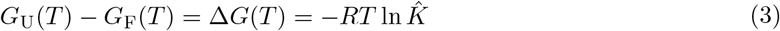

(Barrick, 2018, Chapter 7). From the laws of thermodynamics, the Gibbs free energy of a protein in state X can also be written as *G*_*X*_ = *H*_*X*_ − *TS*_*X*_, where *H*_*X*_ is the molar enthalpy and *S*_*X*_ is the molar entropy of the protein. Enthalpy represents the total heat content of the protein at constant pressure, reflecting the energy stored in its chemical bonds and molecular interactions. Entropy, on the other hand, measures the conformational freedom or disorder of the protein in state X. A folded protein, with fewer conformations, has lower entropy, whereas an unfolded protein, with greater molecular freedom, has higher entropy. Using these definitions, the difference in Gibbs free energy between the folded and unfolded states is given by

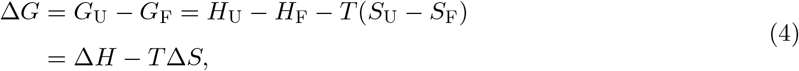

which is valid for any concentrations of folded and unfolded proteins. The change in free energy between the folded and unfolded states is thus equal to the change in enthalpy, which corresponds to the heat absorbed during unfolding, minus the entropy change associated with the conformational changes, scaled by temperature. Combining eqs. (3)-(4) we obtain

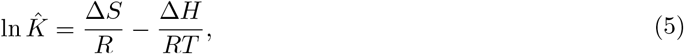

which is known as the Van’t Hoff equation.

### Linear heat capacity model

We now derive the LHC model for the conformational stability of proteins, defined as the free energy difference between folded and unfolded states (eq. 1). In this model, we assume that the heat capacity of a protein in state *X ∈* {U, F} can be expressed as a linear function of temperature,

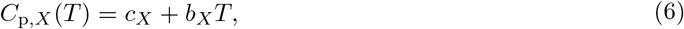

where the subscript “p” denotes that the capacity is defined at constant pressure. The heat capacity *C*_p,*X*_ (*T*) represents the amount of heat required to raise the temperature of a protein in state *X* by one unit, reflecting the system’s ability to store heat energy. The heat capacity is related to the enthalpy of the protein by

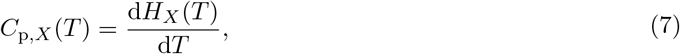

where *C*_p,*X*_ (*T*) is the heat capacity at constant pressure, and *H*_*X*_ (*T*) is the enthalpy of the protein in state *X*, both as functions of temperature. Enthalpy can be expressed as a function of temperature through integration. For a protein in state *X*, this is given by

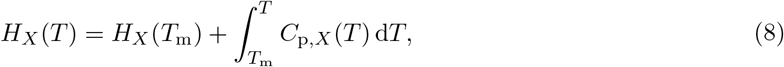

where *T*_m_ is some reference temperature, and *C*_p,*X*_ (*T*) is the heat capacity as defined in eq. (6). Entropy of a protein in state *X* is related to heat capacity as

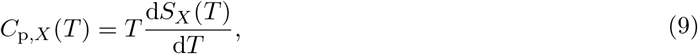

and by integrating we obtain an expression

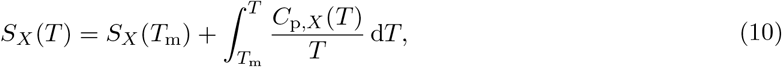

where we used the same reference temperature *T*_m_ as above. Now, by substituting eq. (6) into eqs. (8) and (10), and integrating, the difference in Gibbs free energy (eq. 4) between the folded and unfolded states becomes

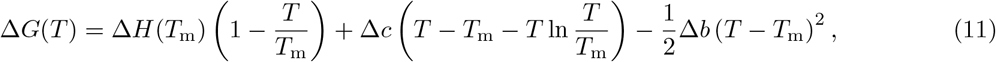

where Δ*H*(*T*_m_) is the difference in enthalpy between folded and unfolded states at *T*_m_, and Δ*c* = *c*_U_ − *c*_F_ and Δ*b* = *b*_U_ −*b*_F_ are the coefficients representing the change in linear heat capacity Δ*C*_p_(*T*) = *C*_p,U_(*T*)− *C*_p,F_(*T*) (eq. 6). This constitutes the linear heat capacity (LHC) model. By setting Δ*b* = 0, we recover the modified Gibbs-Helmholtz equation commonly used in the literature (Becktel and Schellman, 1987; Schellman, 1987; Fersht, 1999; Privalov, 2012; Barrick, 2018). The LHC model can be straightforwardly generalized to accommodate any degree of polynomial.

## 3 Fitting the LHC model to DSC data

Depiction of a typical DSC experiment and the raw DSC output is shown in Figure 1. In a DSC experiment, a solvated protein population in one container and a control solution in a second container, are heated from some initial temperature *T*_min_ at a specific rate *ρ* until the containers reach some final temperature *T*_max_. These temperatures are chosen such that at *T*_min_ the protein population consists of proteins that are mostly in their folded state, and at *T*_max_ where the proteins are mostly unfolded. During the experiment one then keeps track of the difference between the heat capacity of the protein population and the control (y-axis, Figure 1), against the temperature (x-axis, Figure 1), hence the term differential heat capacity and the notation *δC*_p_. The unit of *δC*_p_ is kJ/K/mol or kcal/K/mol.

**Figure 1:**
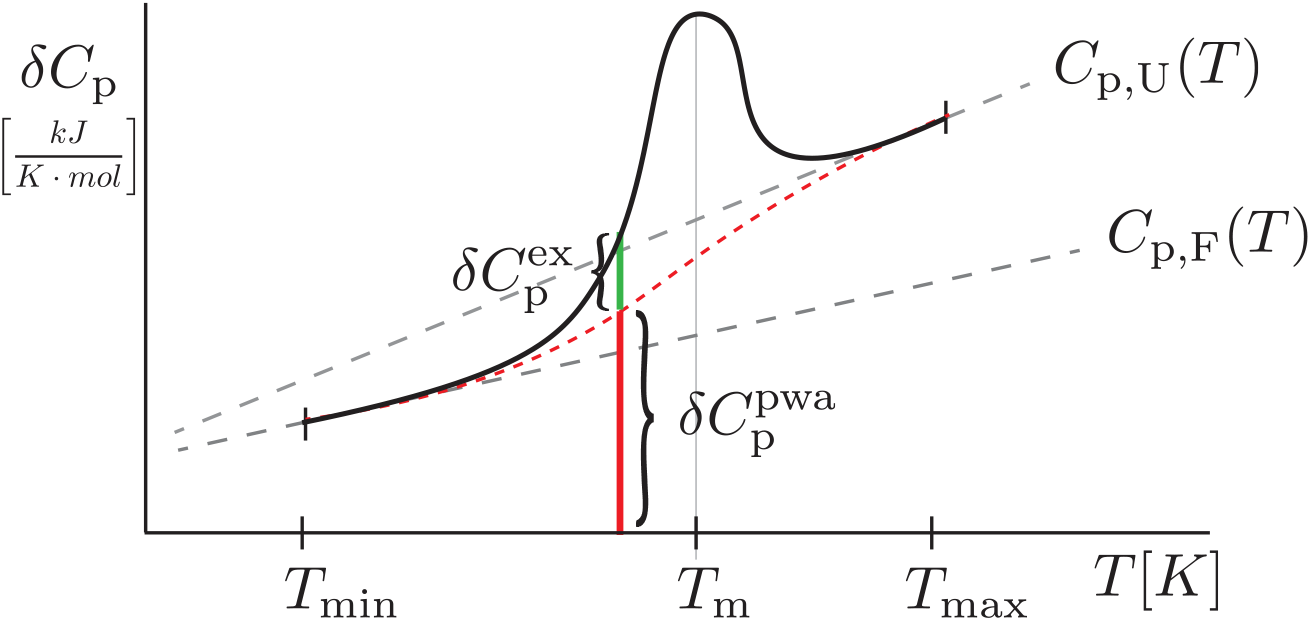
Caricature of raw data from a typical DSC experiment. In the example we assume that the heat capacities of folded and unfolded proteins are linear functions of the temperature (eq. 6), but not necessarily parallel (i.e., in eq. 11 we have Δ*b* ≠ 0). The explanation of the different characters are given in the main text.

The challenge in fitting a model to DSC data arises from the fact that, upon heating, the equilibrium composition of the protein population changes^1^ from being (almost) fully folded at low temperatures to (almost) fully unfolded at high temperatures. This is particularly problematic because the observed differential heat capacity (the thick black concave curve in Figure 1) is not a simple linear combination of the heat capacities of folded and unfolded proteins (the red dotted line in Figure 1). The difference between this linear combination and the observed data is due to proteins undergoing the unfolding transition (green vertical line in Figure 1). This peak results from the unfolding transition and aligns with thermodynamic principles: the energy required to heat the population depends on how the protein population is distributed between the folded and unfolded states (*f*_F_(1 − *f*_F_)). This explains why the peak reaches its maximum when half of the population is folded and the other half is unfolded (because *f*_F_(1 − *f*_F_) is largest when *f*_F_ = 1*/*2). The temperature *T*_m_ at which this occurs is called a melting temperature.

To fit the data, we thus need an expression for the output

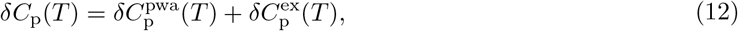

where *δC*_p_(*T*) is the sum of the population-wide average of folded and unfolded differential heat capacities, 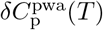, and the excess heat capacity, 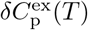 (shown as the vertical red and green lines, respectively, in Figure 1). In other words, we aim to derive an expression for eq. (12) using only a few parameters, which can then be estimated to achieve a good fit to the data. For the population-wide average, we have

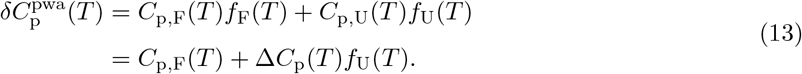

Next, we need an expression for 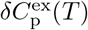. To obtain this, we first derive an expression for the observed differential heat capacity, *δC*_p_(*T*), and then subtract eq. (13). Using the definition of enthalpy (eq. 7), we have

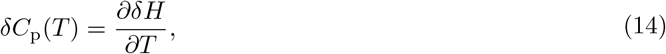

where

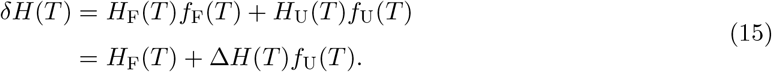

Substituting eq. (15) into eq. (14) and differentiating we get

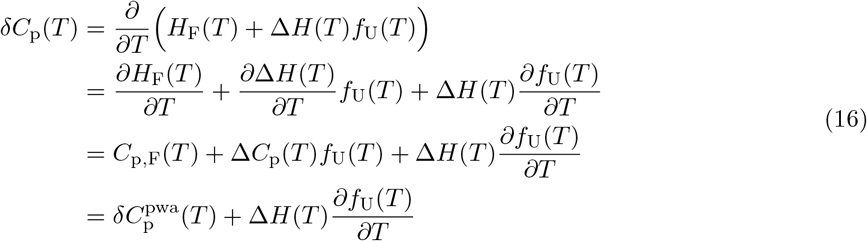

where the third equality follows from eq. (7) and the last equality from eq. (13). Using eq. (12) we get

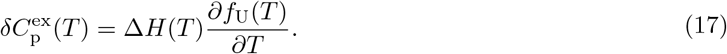

However, since we already need to estimate the parameter *f*_U_(*T*) (which appears in the expression for 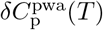 in eq. 13), we aim to avoid introducing an additional parameter, *∂f*_U_(*T*)*/∂T*, especially as it represents the differential of *f*_U_(*T*). The final task is therefore to determine whether we can derive an expression for the differential in eq. (17). This can be done by noting that 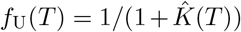 (eq. 2), and hence

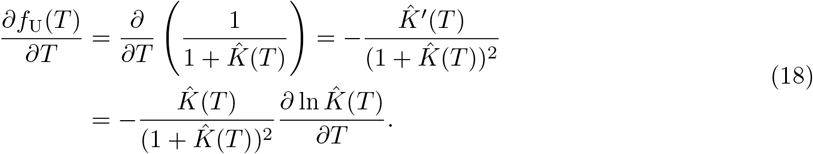

Then, using the Van’t Hoff eq. (5) we have

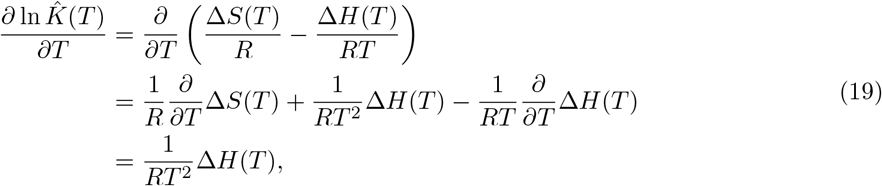

where in the final equality we used the definitions in eqs. (7) and (9). By substituting eq. (19) into eq. (18) we get a relation

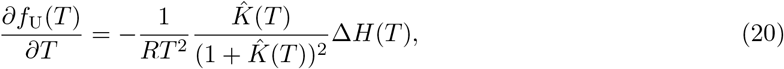

and by substituting this into eq. (17) we finally get

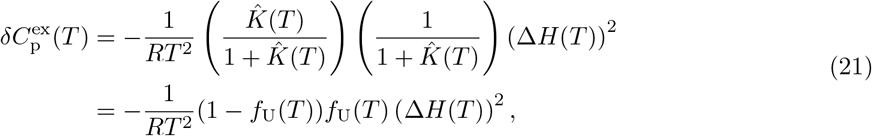

where we again used eq. (2). The expression for the observed differential heat capacity i s thus

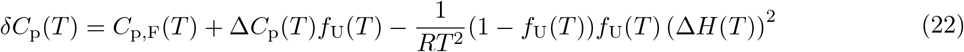

where

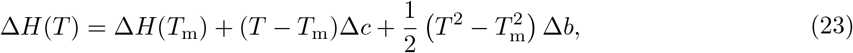

and

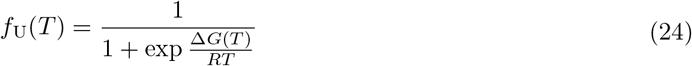

with

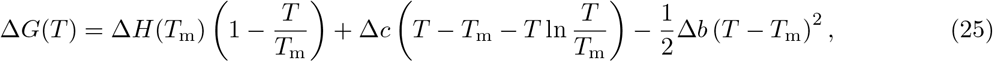

which is just the eq. (11) but we reproduce it here for completeness.

We have thus derived an expression for the differential heat capacity, *δC*_p_(*T*) (eq. 22), where the expression for Δ*H*(*T*) is given in eq. (23) and for *f*_U_(*T*) in eq. (24), into which we substitute an expression for Δ*G*(*T*) as provided in eq. (25). The parameters to be estimated are therefore: the melting temperature *T*_m_, the enthalpy difference at the melting temperature Δ*H*(*T*_m_), the heat capacity of the folded protein *C*_p,F_(*T*), and consequently the parameters *c*_F_ and *b*_F_ (eq. 6), as well as the difference in heat capacity between folded and unfolded proteins, Δ*C*_p_(*T*), leading to the parameters Δ*c* and Δ*b* (eq. 6). To fit the LHC model, we need to estimate 6 parameters, whereas to fit the CHC model, where Δ*b* = 0, we only need to estimate 4 parameters. However, if we assume that the baseline of the experimentally obtained differential heat capacity passes through a specific point derived from the data, we can reduce the number of parameters to be estimated by one. For example, we can get rid of *c*_F_ by setting *c*_F_ = *C*_p,F_(*T*_*◦*_) − *b*_F_*T*_*◦*_ where *T*_*◦*_ and *C*_p,F_(*T*_*◦*_) are obtained from the data. A Mathematica file demonstrating model fitting can be found in the online supplement (Priklopil, 2024).

### 3.1 Examples: fitting protein data

To demonstrate model fitting, we extracted data for the barnase (pH = 2.50) and ubiquitin (pH = 3.0) proteins from the publication by Privalov and Dragan (2007, Fig.1). To analyze this experimental data, the heat capacity plots were converted into a digital format using WebPlotDigitizer (Rohatgi, 2021). We then fitted the LHC and CHC models to the data using Mathematica version 14.0. The two files, one for each protein, are available in the online supplement (Priklopil, 2024), and the results are presented in Figure 2.

**Figure 2:**
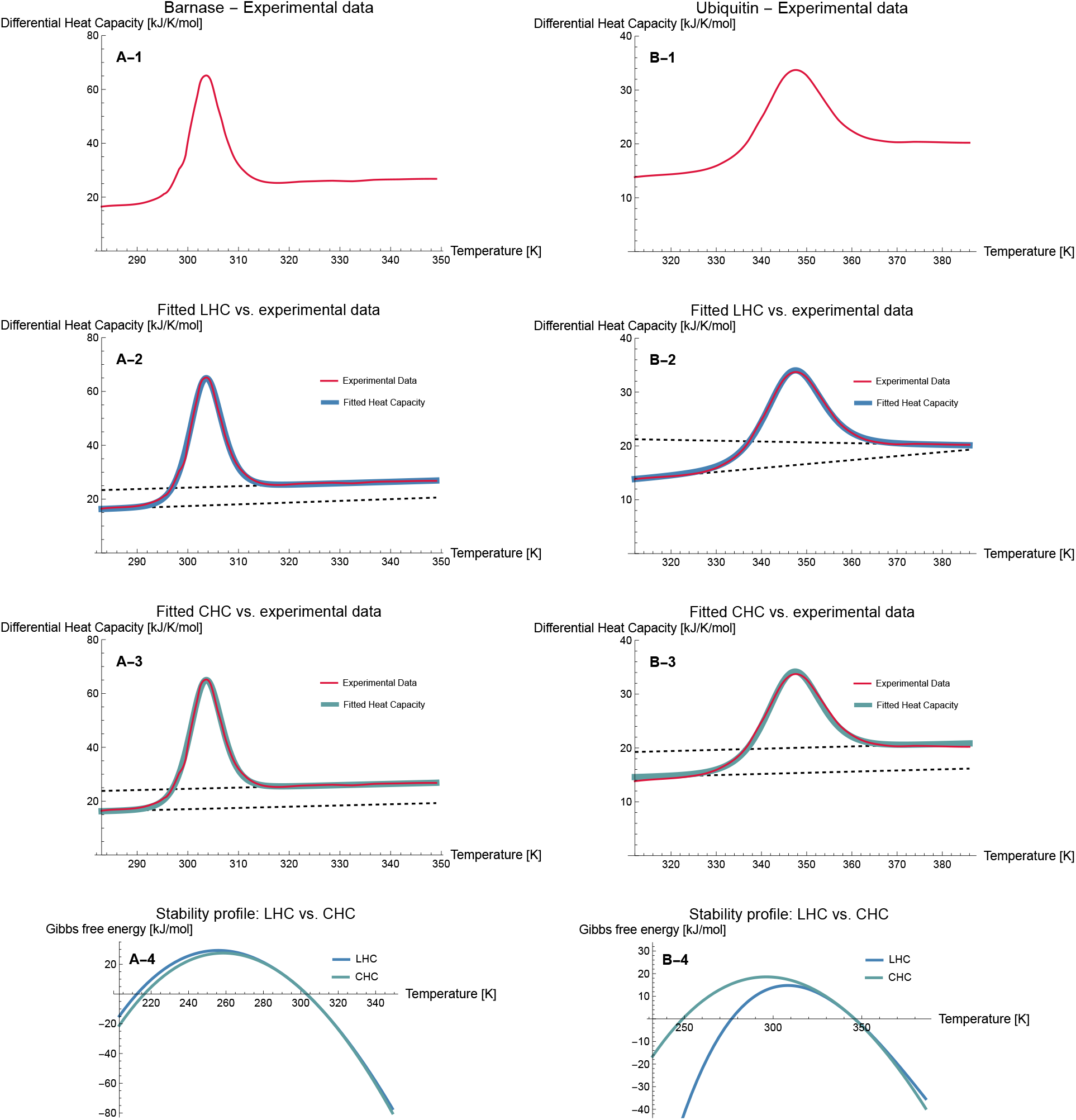
Experimental data and fitted models for Barnase (panels A) and Ubiquitin (panels B) proteins. A-1) Barnase data (pH = 2.50) obtained from Fig.1a in Privalov and Dragan, 2007. A-2) The LHC model was fitted to the data, yielding the following estimates: *T*_m_ = 303.0*K*, Δ*H*(*T*_m_) = 362.4 kJ/mol, *b*_F_ = −0.06 kJ/(mol K), Δ*c* = 10.0 kJ/(mol K), Δ*b* = −0.01 kJ/(mol K). A-3) The CHC model was fitted to the data, yielding the following estimates: *T*_m_ = 303.2 *K*, Δ*H*(*T*_m_) = 364.4 kJ/mol, *c*_F_ = 3.3 kJ/(mol K), *b*_F_ = 0.05 kJ/mol, Δ*c* = Δ*C*_p_ = 7.5 kJ/(mol K) (in this model, Δ*b* = 0). A-4) Using the parameters estimated from panels A-2 and A-3, we plotted the difference in Gibbs free energy (eq. 25) for both fitted models, representing the stability profile of the protein. B-1) Ubiquitin data (pH = 3.0) obtained from Fig.1b in Privalov and Dragan, 2007. B-2) The LHC model was fitted to the data, yielding the following estimates: *T*_m_ = 346.0*K*, Δ*H*(*T*_m_) = 243.7 kJ/mol, *b*_F_ = −0.07 kJ/(mol K), Δ*c* = 35.3 kJ/(mol K), Δ*b* = −0.09 kJ/(mol K). B-3) The CHC model was fitted to the data, yielding the following estimates: *T*_m_ = 345.9 *K*, Δ*H*(*T*_m_) = 251.6 kJ/mol, *c*_F_ = 7.9 kJ/(mol K), *b*_F_ = 0.02 kJ/mol, Δ*c* = Δ*C*_p_ = 4.7 kJ/(mol K) (in this model, Δ*b* = 0). B-4) Using the parameters estimated from panels B-2 and B-3, we plotted the difference in Gibbs free energy (eq. 25) for both fitted models, representing the stability profile of the protein.

In panels A (Figure 2), we analyzed the Barnase data and observed that both models (LHC and CHC) fit well, producing similar stability profiles. This is because the baselines for the folded and unfolded proteins in the data are almost parallel, resulting in comparable fits for both models. In contrast, in panels B (Figure 2), where we analyzed the Ubiquitin data, the baselines are not parallel. Consequently, the data does not align well with the constant heat capacity model, which assumes parallel baselines. As a result, the estimated parameters lead to notably different stability profiles (panel B-4, Figure 2). We also note that the estimates for the CHC model are similar to those obtained by Yeritsyan and Badasyan (2024a, Barnase: *T*_m_ = 303 *K*, Δ*H*(*T*_m_) = 364 kJ/mol, Δ*C*_p_ = 5.6 kJ/(mol K); Ubiquitin: *T*_m_ = 346 *K*, Δ*H*(*T*_m_) = 254.5 kJ/mol, Δ*C*_p_ = 3.85 kJ/(mol K)), although it remains unclear how the baseline was handled in their model fitting.

## 4 Conclusions

In this note, we derived a linear heat capacity model step-by-step to facilitate the analysis of thermodynamic parameters of two-state proteins using data from differential scanning calorimetry experiments. While a systematic comparison of our model’s goodness-of-fit relative to previous models is beyond the scope of this work, our analysis of two proteins suggests that this model can provide an improved fit, warranting further research. This work has potential applications across fields, from protein engineering to studies of natural populations, and we hope it proves valuable to readers interested in the mathematical foundations of protein stability analysis.

## Footnotes

1 Recall that we assume that the heating process is much slower than the folding and unfolding of proteins, allowing them to be in equilibrium at all times.

